# Subthalamic beta bursts correlate with dopamine-dependent motor symptoms in 106 Parkinson’s patients

**DOI:** 10.1101/2022.05.06.490913

**Authors:** Roxanne Lofredi, Liana Okudzhava, Friederike Irmen, Christof Brücke, Julius Huebl, Joachim K. Krauss, Gerd-Helge Schneider, Katharina Faust, Wolf-Julian Neumann, Andrea A. Kühn

## Abstract

Pathologically increased beta power has been described as a biomarker for Parkinson’s disease (PD) and related to prolonged bursts of subthalamic beta synchronization. Here, we investigate the association between subthalamic beta dynamics and motor impairment in a cohort of 106 Parkinson’s patients in the ON- and OFF-medication state, suing two different methods of beta burst determination. We report a frequency-specific correlation of low beta power and burst duration with motor impairment OFF dopaminergic medication. Furthermore, reduction of power and burst duration correlated significantly with symptom alleviation through dopaminergic medication. Importantly, qualitatively similar results were yielded with two different methods of beta burst definition. Our findings validate the robustness of previous results on pathological changes in subcortical oscillations both in the frequency-as well as in the time-domain in the largest cohort of PD patients to date with important implications for next-generation adaptive deep brain stimulation control algorithms.

## Introduction

Intracerebral recordings from patients undergoing deep brain stimulation (DBS) surgery for Parkinson’ disease (PD) have revealed increased subthalamic oscillatory beta band activity (13-35 Hz) as a pathophysiological hallmark of the PD motor state^1–6^. Both dopaminergic medication as well as DBS are associated with a decrease of this pathologically enhanced activity, specifically in the low beta sub-band (LB=13-20 Hz), which has primarily been related to bradykinetic-rigid symptoms^7–12^. However, amplitude and temporal dynamics of high beta activity (20-35 Hz) seems implicated in bradykinetic-rigid symptoms^13,14^ and gait difficulties in PD as well^15,16^. Taken together, subthalamic beta activity has been shown to correlate with the entire spectrum of motor symptoms assessed by the Unified Parkinson’s Disease Ratings Scale (UPDRS) in the dopamine-depleted state^9,17^, even months after chronic stimulation^18,19^. Thus, beta activity is considered a stable biomarker of motor signs in PD and has been used as feedback signal for therapeutic demand in experimental studies on adaptive closed-loop DBS (aDBS) with similar or better efficacy / side-effect profile than conventional DBS ^20–23^.

The beneficial effects of aDBS have been linked to a shortening of pathologically prolonged periods of beta synchronization, so-called beta bursts. In contrast, the overall decrease of beta burst duration with conventional DBS^24^ possibly relies on suppression of both short and long beta bursts ^25,26^. Suppressing short beta bursts, however, might be disadvantageous as they are considered part of physiological signaling within the motor circuit^27^. Accordingly, a shift towards shorter beta bursts is observed with dopaminergic medication^2,12,13^, which has been correlated with clinical improvement in one study so far. With increasing interest for the pathophysiological role of beta bursts in PD, novel methods of burst definition have recently been proposed ^28^. These have been validated by investigating DBS-induced changes but not used for capturing dopamine-related dynamics of beta bursts yet ^24,29^.

Most studies on oscillatory activity of DBS-target structures have been conducted during or in the days following the implantation of DBS-electrodes, where the leads are still externalized, as chronic recordings from implanted pulse generator have become available only in the last couple of years^30–33^. Subcortical spectral densities and local cross-frequency coupling in the dopamine-depleted state of PD-patients have been assessed in large cohorts of >50 PD patients (n=63 in Neumann et al., 2016; n=74 in Shreve et al., 2017)^17,34^. However, given that the dopamine-substituted state can rarely been assessed in the intra-operative setting and the concept of the pathophysiological role of temporal dynamics in oscillatory activity in PD emerged only recently, studies on dopaminergic effects on beta power or beta burst dynamics rely on much smaller cohorts of <25 PD patients^9,12–14,24,35^.

The present study aims to test the robustness of previous results on a dopamine-dependent correlation between subthalamic beta burst dynamics and motor symptoms in Parkinson’s disease by i) validating these findings in the largest cohort to date of > 100 PD patients and ii) comparing an established with a newly developed method of beta burst assessment.

## Results

### Correlation of beta power and motor impairment

Relative power spectra were averaged across contact pairs separately for the ON- and OFF-medication state. Each frequency bin was tested for difference across medication states, revealing a significant decrease (P<0.001) of relative power between 10-19 Hz with dopaminergic medication when averaged across all channels or 10-20 Hz when only considering the channel with most pronounced LB peak per hemisphere in the OFF medication state (see Figure 1A). While significant correlation with symptom severity in the OFF medication state was found in a slightly broader spectrum (9-22 Hz), only frequencies within the LB band (13-19 Hz) were significantly associated to symptom alleviation with dopaminergic medication (see Figure 1B). To remain comparable with previous studies, the following analyses were performed with a predefined LB band from 13-20 Hz, but results remained stable if the following analyses were performed by averaging over 13-19 Hz. When averaged across the LB band, there was a significant correlation between motor impairment as assessed by the UPDRS-III and LB power in the OFF-medication state (Rho=0.21, P=0.03, see Figure 1C). While this correlation was not significant in the ON-medication state (P>0.05), the dopamine-related motor improvement (UPDRS OFF-ON) was significantly associated with the reduction in LB band power (Rho=0.36, P=0.0008, see Figure 1C) as well. Clinical subtypes showed no differences in overall LB power (P>0.05). However, there was a symptom-specific correlation between LB power in the OFF medication state with bradykinesia (Rho=0.3, P=0.04) but not rigidity or tremor in the subset of patients in which UPDRS subscores were available (see Supp. Fig. 1). In contrast, there was no symptom specific correlation with LB power in the ON medication state, nor with medication-related symptom alleviation. Overall, there was no significant correlation of motor impairment or improvement with averaged HB power or averaged alpha power in none of the tested conditions mentioned above. Spectral peaks in the LB and HB band showed a similar distribution across contact pairs, with most peaks found in the more dorsal contact pairs 23 and 12 (LB: 82% and HB: 83% of all detected peaks). Distribution of alpha band peaks showed a more variable distribution across contacts with about 30% of detected peaks found in the lowermost contacts, see Supp. Figure 2. For a more detailed analysis of peak localization in this patient cohort, see Darcy et al., 2022^36^.

**Figure 1:**
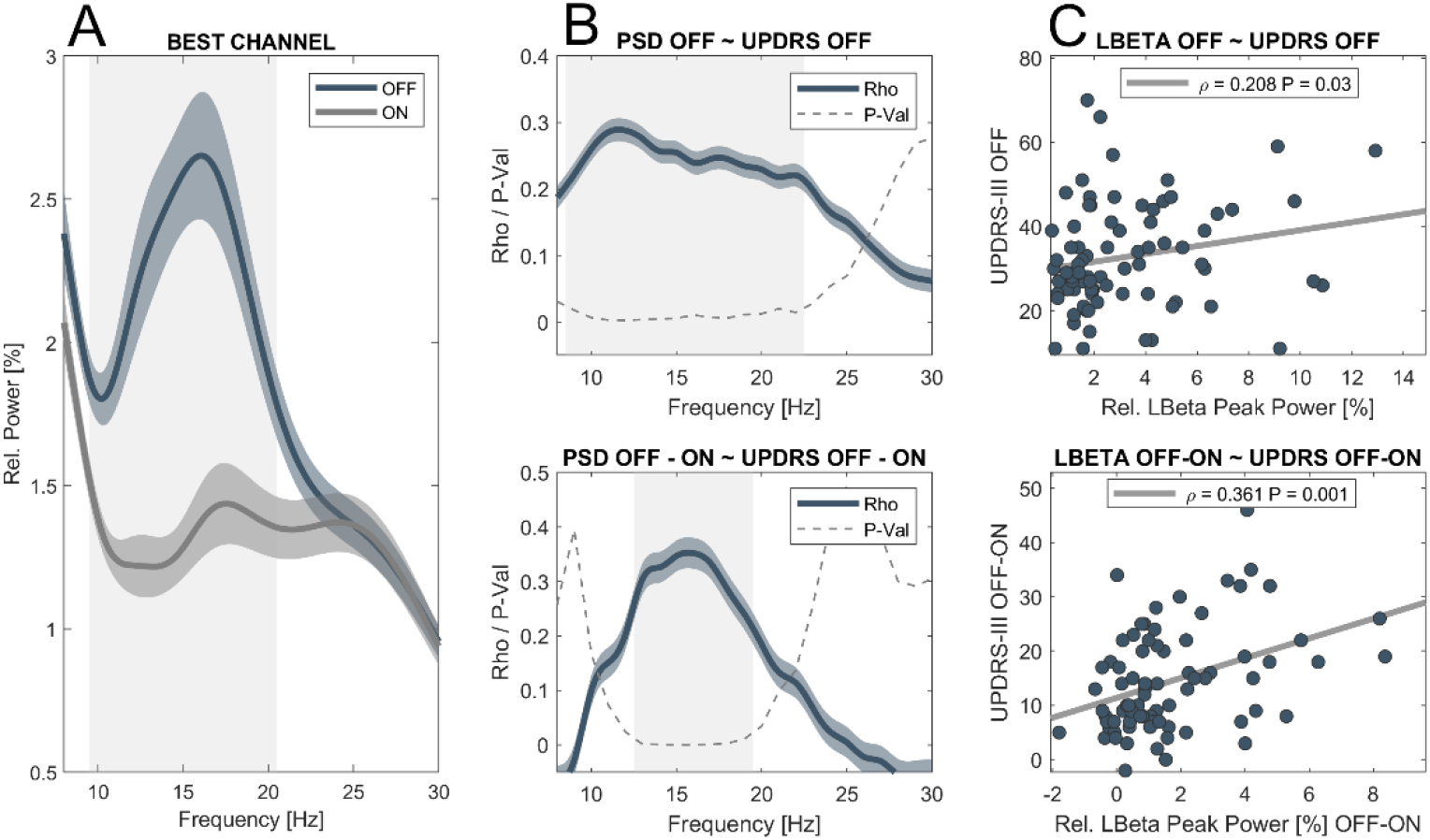
Low beta power in the OFF-medication state and its reduction with dopamine correlates with symptom severity. (A) Averaged power spectra across contact pairs in the ON (grey) and OFF (blue) medication state. Note the decrease in relative power with dopaminergic medication that includes the low beta and sub-beta band (underlined in grey, P<0.0001). (B) Matching the frequency bins that show a significant modulation with dopamine, amplitude of frequencies between 9-22 Hz show a significant correlation with motor symptoms as assessed by the UPDRS-III. However, symptom alleviation with dopamine is best reflected by amplitude changes from 13-19 Hz, commonly referred to as low beta band. In blue are shown Rho-values for each frequency bin, the scattered grey line shows the correspondent P-value, significant areas are underlined in grey. (C) There is a significant correlation between symptom severity and averaged low beta power (13-20 Hz) in the dopamine-depleted state (Rho=0.208, P=0.03). Likewise, the reduction in low beta power with dopamine correlates with symptom alleviation (Rho=0.36, P=0.0084, right column).

### Correlation of beta burst duration and motor impairment

When using a baseline as established in Anderson et al., 2020, averaged LB burst duration was significantly prolonged after 12-hour withdrawal of dopaminergic medication (OFF=579.1± 469.1 ms; ON=358.6 ± 230.2 ms; P<0.0001), see Figure 2A. In the OFF-medication state, the averaged LB burst duration correlated significantly with motor impairment (Rho=0.18, P=0.05) as did the shortening of mean LB burst duration with motor improvement (Rho=0.26, P=0.008), see Figure 2B. There was a symptom-specific correlation of LB burst duration in the OFF medication state with bradykinesia (R=0.31, P=0.03) and rigidity (R=0.35, P=0.01) but not tremor symptoms in the subset of patients with UPDRS subscores (see Supp. Fig. 1). There was no significant correlation between LB burst duration and motor impairment in the ON-medication state. When burst time bins were tested separately, motor impairment in the OFF state was positively correlated with the amount of bursts above 500 ms length (500-600 ms: Rho=0.29, P=0.005; 600-700 ms: Rho=0.26, P=0.01) and negatively correlated with short beta bursts (100-200 ms: Rho=-0.18, P=0.05; 200-300 ms: Rho=-0.2, P=0.03), see Figure 2C. Accordingly, changes in short and long bursts may best reflect symptom alleviation with medication (100-200 ms: Rho=-0.19, P=0.04; 200-300 ms: Rho=-0.2, P=0.03; > 700 ms: Rho=0.26, P=0.02), see Fig.2C. However, these time-bin specific results did not survive FDR-correction for multiple comparisons. There was no significant correlation between motor symptoms and LB burst amplitude or HB burst properties in either medication state.

**Figure 2:**
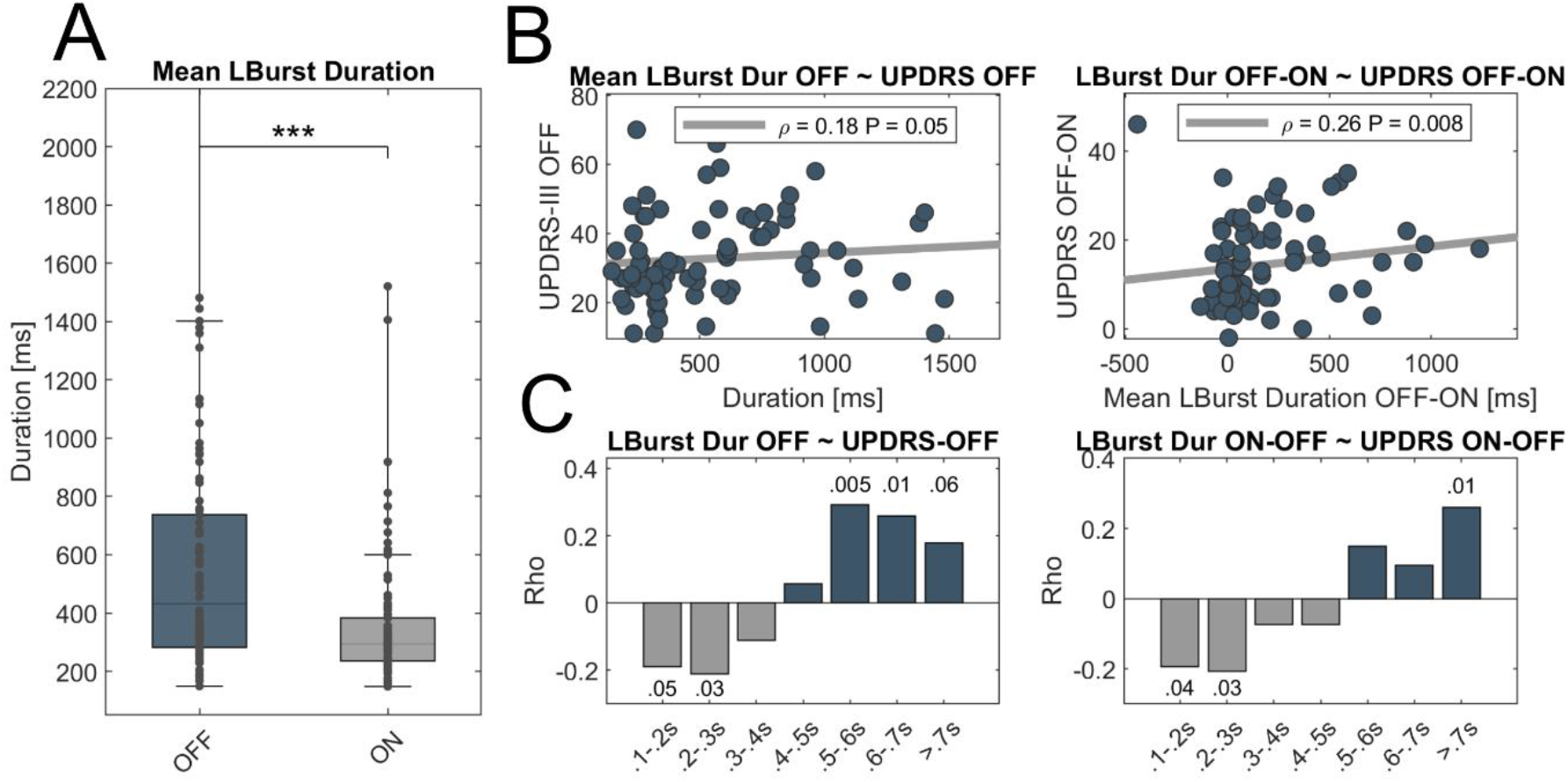
Low beta burst duration decreases with dopamine and correlates with symptom severity. (A) There is a significant shortening of averaged low beta burst duration in the ON-medication state (P<0.001). (B) When averaged, the mean LB burst duration correlates with motor impairment in the OFF medication state (left column, Rho=0.18, P=0.05) and symptom alleviation with medication (right column, Rho=0.26, P=0.008). (C) While the amount of short LB bursts correlates negatively, the amount of long LB bursts correlates positively with motor impairment (left column). A similar distribution is seen for the correlation with dopamine-related symptom alleviation and change in amount of short vs long LB bursts (right column). However, the time-bin related results do not remain significant after FDR-correction. Shown are P-values per time-bin before FDR-correction. In boxplots, central marks indicate the median and edges the 25th and 75th percentiles of the distribution. *** p<0.001.

### Comparison of different methods for burst detection

As expected, beta bursts where significantly shorter when using the common threshold method based on the 75th percentile of the amplitude distribution (OFF=304.9± 42.3 ms vs 579.1± 469.1 ms, P<0.001; ON=289.4 ± 36.7 ms vs 358.6 ± 230.2 ms; P=0.001). However, there was a significant correlation between averaged burst duration between both methods (OFF: Rho=0.56, P<0.001; ON: Rho=0.63, P<0.001), specifically with regard to the amount of very short and very long LB bursts in both medication states (OFF: 100-200 ms: Rho=0.46, P<0.001; >700 ms: Rho=0.53, P<0.001; ON: 100-200 ms: Rho=0.52, P<0.001; >700 ms: Rho=0.53, P<0.001).

In the OFF medication state, common threshold based results were qualitatively similar to the baseline-based results with a frequency-specific correlation between mean LB burst duration and motor impairment (Rho=0.27, P=0.008) that may rely on the amount of long bursts (>700 ms: Rho=0.22, P=0.02) and negatively associated with the amount of very short bursts (100-200 ms: Rho=0.18, P=0.05; uncorrected). However, dopamine-related changes were less well captured by the common threshold based method. While a significant reduction of averaged burst duration with medication was observed (OFF=304.9± 42.3 ms; ON=289.4 ± 36.7 ms; P<0.0001), its amount did not correlate with symptom alleviation, neither when averaged across all bursts nor when considered separately for each time bin. For a summary of common threshold based results, see Supp. Fig 3.

## Discussion

In the present study, we investigated the association between subthalamic beta power and bursts with motor symptoms as well as their change with dopaminergic medication in a large cohort of 106 PD-patients. We show a significant correlation between motor symptom severity in the OFF medication state and oscillatory activity from ~10-20 Hz. Moreover, the medication-related reduction of LB power (13-20 Hz) was paralleled by a dopamine-related symptom alleviation. Similar to LB power, averaged LB burst duration significantly correlated with motor impairment in the OFF medication state, as did dopamine-related reduction of LB burst duration with symptom alleviation, when using the noise floor as baseline. All observed effects were frequency-specific for the LB band and a sub-analysis suggested that they might be driven by bradykinetic symptoms. We thus provide further evidence for temporal dynamics of LB activity as a robust biomarker of motor symptoms in a large cohort of PD patients, relatively independent of the exact method used for burst determination.

### Frequency-specific prolongation of low beta bursts in the dopamine-depleted state

Our results are in line with previous studies, demonstrating a correlation of parkinsonian symptoms in the OFF medication state^17^ and a specific dopamine-sensitivity of activity mainly covering the LB band^37^, which has also been described for DBS-related effects^10^. A recent study combining magnetoencephalography, subthalamic vs. pallidal LFP and computational modelling, has attributed the difference of high- and low-beta to the hyperdirect- and indirect-pathway^38^. The findings support the notion that PD symptoms may arise from hyperactivity of the indirect pathway, while hyperdirect cortico-subthalamic communication is implicated in cognitive control and motor inhibition^39,40^.

Regarding the more recently emerged concept of pathologically prolonged beta bursts in the OFF medication state, frequency-specificity for the LB band has been explicitly reported only in one study investigating the association between beta bursts and bradykinesia^11^. In two other studies, beta bursts were determined around beta peak frequencies, the average of which lied in the LB band as well (Tinkhauser et al., 2017: 19.9 Hz; Lofredi et al., 2019: 14.3 Hz). Not all studies have produced this distinction of high- and low beta activity, e.g. a more recently published study reported a frequency-specific correlation of high beta burst duration with motor symptoms^13^. The precise differences remain unknown, but it is important to note that potentially electrode impedance and spatial coverage may have an impact on the recorded frequency bands in LFP^41^. All of the aforementioned studies however reported results on <30 PD-patients. The present study provides further evidence that in a large cohort, the best across patient correlation with symptom severity is obtained with burst properties in the LB band.

### Qualitatively similar results with different methods for burst determination

Methodologically, a vivid scientific controversy has risen around the potential pitfalls of burst determination. Earlier studies established thresholds for bursts based on amplitude distribution either by considering only amplitudes above a specific percentile^12^ or median deviation ^42^ that were applied by following studies ^2,11,13,43–45^. Others tried to train discriminators on artificially generated time series for burst detection, which is a promising direction, but might need further knowledge on the underlying amplitude distribution before assuring to generate physiologically meaningful results^25^. More recently, a novel method of burst determination, using a physiological baseline based on the recording-specific noise floor was proposed ^28^ and used in several studies investigating DBS-associated beta burst dynamics in PD-patients with chronically implanted pulse generators^24,29^. However, this method had not been used to assess dopamine-related beta burst dynamics in externalized recordings of PD patients yet. Here, we obtain similar burst durations in the OFF medication state than these studies, that were moreover qualitatively similar, in both used methodological approaches. This provides further evidence for a stable, hardware-independent, medication-as well as stimulation sensitive and thus pathophysiological relevant correlation between temporal dynamics of LB activity and motor state in PD patients.

### Limitations of beta power and beta bursts as a biomarker

Although subthalamic LB power and burst duration appear as robust biomarkers across PD-patients, the explained variance of motor symptom severity remains relatively low, with rho-values ranging around 0.4-0.5 in previous studies and 0.18-0.36 in the present study with the largest cohort to date. This low explanatory power might relate to the following aspects: First, subthalamic LB power and burst duration might reflect similar aspects of the parkinsonian motor state as burst duration and amplitude are highly correlated. Thus, combining both does not significantly increase the explained variance. Second, LB power and burst properties lose their explanatory value for symptom severity when considering the ON medication state and additional features beyond beta, such as movement- or dyskinesia-related low-frequency or gamma activity may be more performative in this state^43,46^. Third, an individualized investigation of combined spectral changes assessed on a case-by-case basis avoiding the need for spectral normalization, may outperform the information content of spectral features based on group results for individual patients. Thus, while our study provides clear evidence for the pathophysiological relevance of LB in PD, other frequency bands may be better suited to act as control signals in individual patients. Also, cortical signals might be advantageous when compared to subcortical activity patterns in decoding specific motor states^47,48^. In the same vein, multisite recordings could be necessary if the entirety of motor- and non-motor symptoms have to be captured, given that PD is considered a network disease and pathological interregional connections have been described, specifically for motor cortical areas^49–51^. Interestingly, a first pioneering study investigating symptom correlations from combined chronic ECoG and LFP recordings in freely moving patients found different combinations of recording source (Motor cortex vs. STN vs. Motor cortex – STN coupling) and frequency band (alpha, beta, gamma) for each of the five patients to be most predictive of symptom severity^52^.

### Outlook

In conclusion, our study corroborates and extends previous findings on the pathophysiological relevance of beta burst dynamics in PD. Moreover, our findings highlight the importance of understanding temporal brain circuit dynamics to inspire the design of bidirectional control algorithms for on-demand adaptive neuromodulation. The fact that the shift from very short to pathologically prolonged bursts can be observed in such a large cohort of PD-patients increases the confidence that an aDBS-approach, where stimulation would be triggered by beta amplitude threshold crossings of specific lengths, can be viable and advantageous in clinical settings. Large multicenter trials are currently in development with manufacturers offering first generations of commercially available sensing enabled devices. Our results suggest that aDBS trials should implement temporal threshold parameters beyond the single or dual power threshold paradigms to consider temporal burst dynamics as a key hallmark of pathological basal ganglia activity in PD.

## Methods and Materials

### Patients and Surgery

For this study, we have identified archival local field potential (LFP) data from 106 PD-patients (63.5±8 years, 42 female, clinical subtype: 47% akinetic-rigid; 20% tremor-dominant; 33% equivalent) who underwent bilateral implantation of subthalamic DBS-electrodes. For detailed clinical information, see Table 1. In 95 cases, assessment of motor impairment using the UPDRS-III (denominated UDPRS in the following) was available both under regular dopaminergic medication (ON state) and at least 12 hours after dopaminergic drug withdrawal (OFF state) either pre-operatively (n=28) or during the recording sessions (n=67). In a subset of patients (n=45), sub-scores for bradykinesia (UPDRS III items 6.a-9.b, item 14), rigidity (UPDRS-III items 5.a-5.e) and tremor (UPDRS-III items 3.a-4.b) were available in both medication states and used for a symptom-specific sub-analysis of neurophysiological markers. Motor impairment was significantly more pronounced in the OFF than in the ON medication state (UPDRS-OFF: 32.7±12.7, UPDRS-ON: 18.3±9.3, P=0.002). All patients gave written informed consent, which was approved by the local ethics committees (Charité – Universitätsmedizin Berlin and Medizinische Hochschule Hannover) in accordance with the standards set by the Declaration of Helsinki. Three types of DBS-macroelectrodes were used: Medtronic 3389 (n=80), Boston Vercise with circular electrodes (n=11) and Boston Vercise Cartesia (n=15), in which mid-contacts are segmented. Intraoperative microelectrode recordings, test stimulation and postoperative imaging ensured correct DBS-electrode placement. A subset from a previous study (n=45) that assessed beta power in the OFF medication state of PD patients have been included in the present cohort^17^.

### Recordings

Subthalamic LFP recordings of at least 200 seconds were performed in the ON- and OFF medication state in the 1-7 days following DBS-surgery. Patients were comfortably seated and instructed not to move during the recording. Signals were amplified (50.000x) using a D360 amplifier (Digitimer, Hertfordshire, UK) and recorded at a sampling frequency of 1 kHz through a 1401 A-D converter (CED, Cambridge, UK) onto a computer using Spike2 software. When electrode contacts were circular, bipolar LFPs were recorded between adjacent contact pairs. In directional leads, recordings were referenced to the lowermost contact of the electrode, subsequently summed and re-referenced offline to approximate the bipolar recordings derived from adjacent pairs of circular contacts, for example: (contact 1/2 + contact 1/3 + contact 1/4)-(contact 1/5 + contact 1/6 + contact1/7).

### Signal Processing

Artifact-free recording segments were identified visually and analyzed using custom MATLAB code (The Mathworks, Natick, Massachusetts) based on SPM12 (http://www.fil.ion.ucl.ac.uk/spm/) and FieldTrip (http://fieldtrip.fcdonders.nl/). Continuous recordings were filtered (high pass=5 Hz, low pass=98 Hz and notch filter=48-52 Hz) and transferred into the frequency domain using Morlet wavelets with 10 cycles and a temporal resolution of 200 Hz. Power spectra were derived through averaging over time and normalized to the total sum of the recording and further expressed as a percentage of total power. Normalization allows comparison across participants as arbitrary amplitude differences due to electrode impedance, specifically related to the surface difference in segmented vs ring electrodes, and distance to source are removed. Frequency bands of interest in this study where predefined as alpha (8-12 Hz), low beta (LB, 13-20 Hz) and high beta (HB, 21-35 Hz) band. Spectral peak frequencies and the channel displaying the highest spectral peak in the respective frequency bands of interest (alpha=8-12 Hz, LB=13-20 Hz, HB=21-35 Hz) were automatically detected, see Figure 3. For correlative analyses with motor impairment, power values in each frequency band were calculated for the channel with highest peak frequency amplitude in the OFF medication state, separately for each hemisphere and averaged across hemispheres. There was no significant shift in peak frequency from the OFF to the ON medication state (P>0.05) and no difference in band specific power between male and female subjects (P>0.05) in either frequency band.

**Figure 3:**
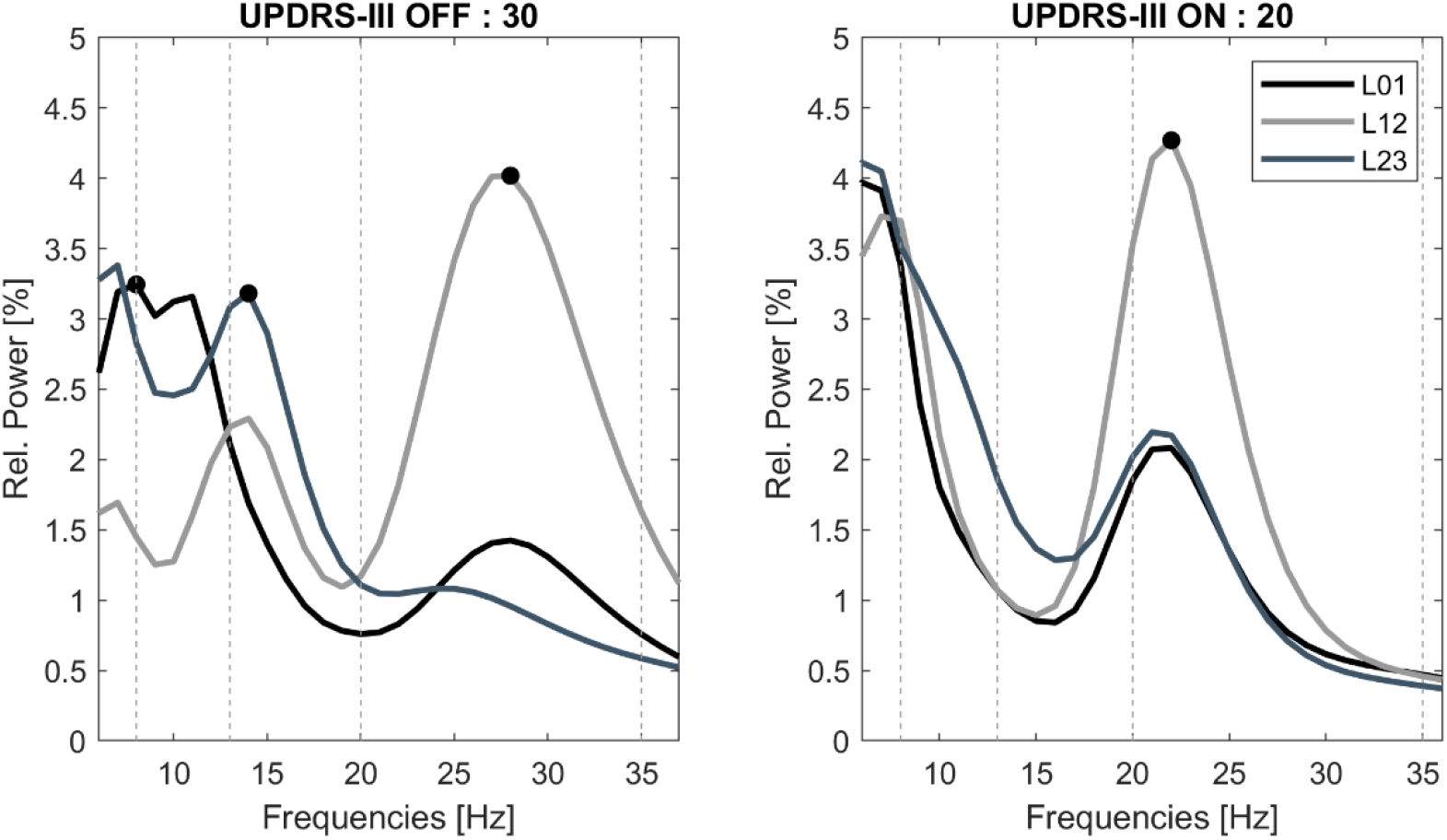
Exemplary trace of power spectra in the ON- and OFF-medication state. Note the multiple spectral peaks can be detected within the same patient or even the same recording channel in the alpha, low and high beta band. Specifically the detected peaks in the low beta range decrease in the ON medication state, while high beta activity remains prominent. Peak frequency with highest amplitude is labeled with a black dot, separately for each respective frequency band of interest. Shown is the left hemisphere of subject 31.

### Beta Burst Analysis

For burst determination, the bipolar channel with highest peak in the respective frequency band was selected in the OFF medication state and used for both medication states. Two methods were used separately for burst determination and compared qualitatively. The first method was recently established and qualifies beta bursts when beta activity exceeds the overall noise floor within a recording ^24,28^. For this approach, the raw signal was bandpass-filtered around the frequency band of interest (here low or high beta band) and envelope peaks of the squared signal were linearly connected to generate an amplitude envelope. A beta burst was detected when the amplitude crossed a threshold defined as four times the median of averaged troughs from envelopes of five overlapping 6 Hz bands in the low gamma band (55-65 Hz) from the same recording. The second method was used in previous publications by the authors^2,11,53^, qualifiying beta bursts as events in which beta activity is signifcantly higher than the overall beta amplitude distribution. Here, the wavelet amplitude was separately averaged across the respective frequency band of interest and smoothed with a moving average Gaussian smoothing kernel (175 ms). Each power amplitude trace was z-scored (X-μ/δ) over the entire recording, separately for the ON- and OFF-medication state. The threshold for bursts was defined as the 75th percentile of the normalized signal amplitude distribution. A separate threshold was calculated in the ON and OFF medication state and the average of both was applied for burst determination in both medication states. In both methods, beta burst duration was defined as time spent above the defined threshold. The distribution of burst durations was considered by categorization into 7 time windows of 100 ms starting from 100 ms to > 700 ms in duration. As for power analyses, obtained values were averaged across hemispheres for correlative analyses with motor impairment.

### Statistics

First, each frequency bin (1 Hz) of the power spectral density was tested for i) changes with medication and ii) correlation with symptom severity as assessed by the UPDRS-III. In the following, power was averaged over canonical frequency bands, low (13-20 Hz) and high (21-35 Hz) beta band respectively, and tested for correlation with symptom severity. For burst analyses, averaged burst duration as well as relative amount of bursts with duration between 100 ms and >700 ms were correlated with UPDRS-III scores. In a sub-analysis with 49 PD-patients, averaged power and burst duration was separately correlated with summed scores for bradykinesia, rigidity and tremor. Across all analyses, nonparametric Monte Carlo permutation tests with 5000 permutations were used for statistical analyses as they do not rely on assumptions about the underlying data distribution. For correlative analyses, Spearman’s correlation (denoted as Rho) was calculated and tested for significance using permutation. All results are indicated as mean ± standard deviation and reported significant at an α level of 0.05. Multiple comparisons were controlled by using detection of the false discovery rate (FDR) where appropriate.

### Data availability statement

The data that support the findings of this study are available on request from the corresponding author in the framework of a data sharing agreement. The data are not publicly available as this would compromise the privacy of research participants according to the current General Data Protection Regulation of the European Union.

## Supp. Figures

**Supp. Figure 1:**
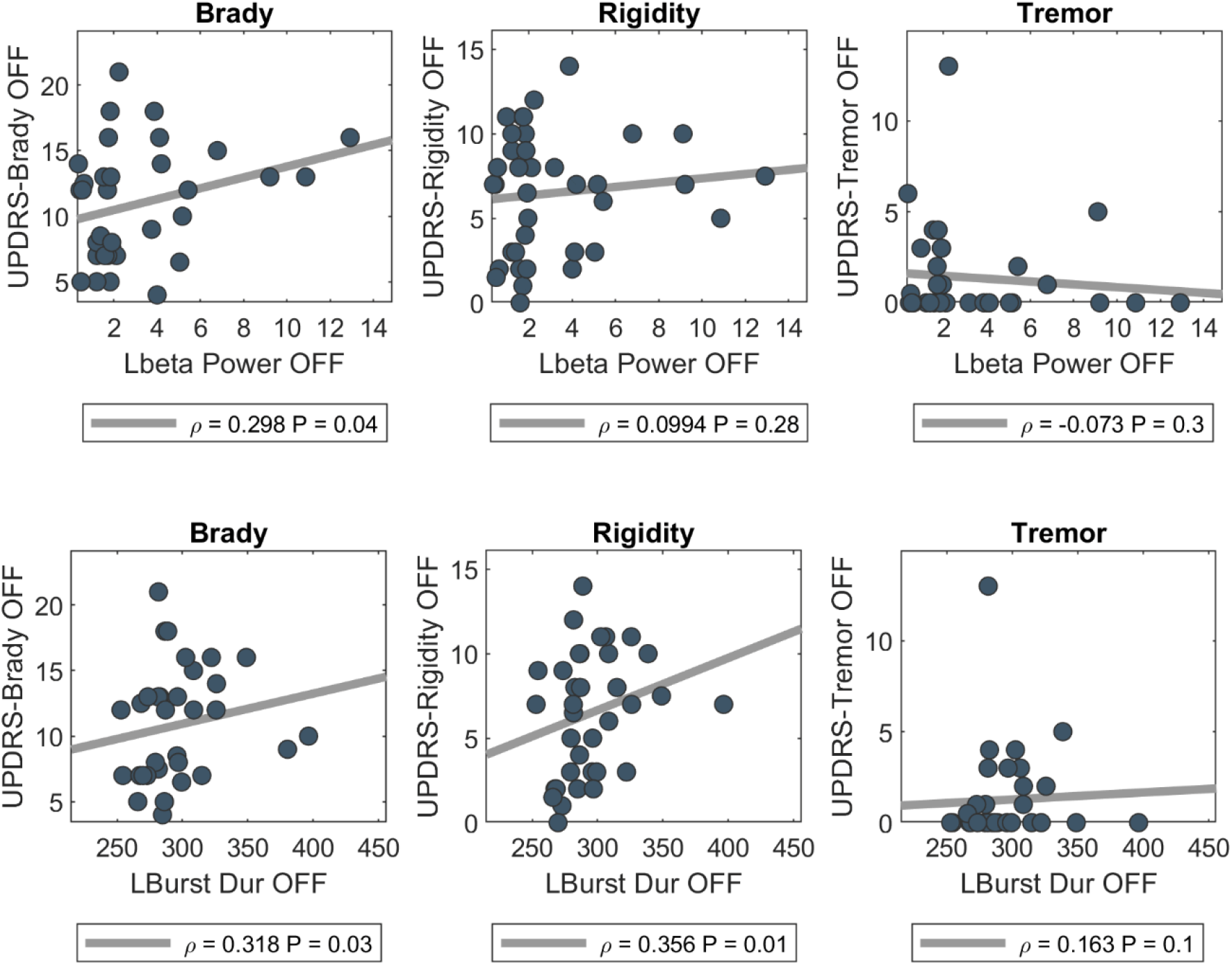
LB power and burst duration as symptoms specific marker. Shown are spearman’s correlation coefficients of LB power (upper row) and LB burst duration (lower row) in the OFF medication state with summed scores for the UPDRS-items bradykinesia (item 6.a-9.b, left column), rigidity (item 5, mid column) and tremor (item 3.a-4.b, right column). Not FDR-corrected.

**Supp. Figure 2:**
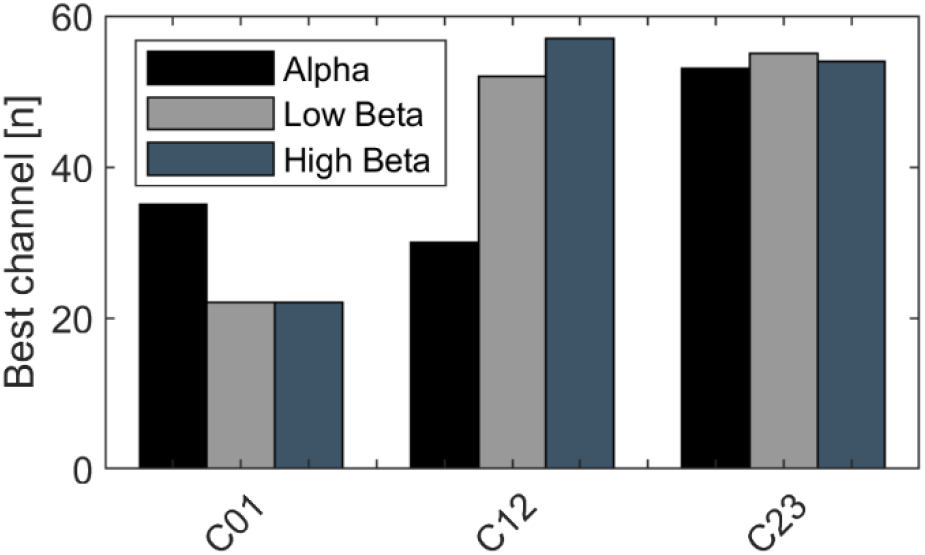
Distribution of spectral peaks across contact pairs. Shown is the number of cases in which the most pronounced spectral peak of the respective frequency band (alpha, low beta and high beta) occurred in the lowermost (C01), mid (C21) or uppermost (C23) contact pair.

**Supp. Figure 3:**
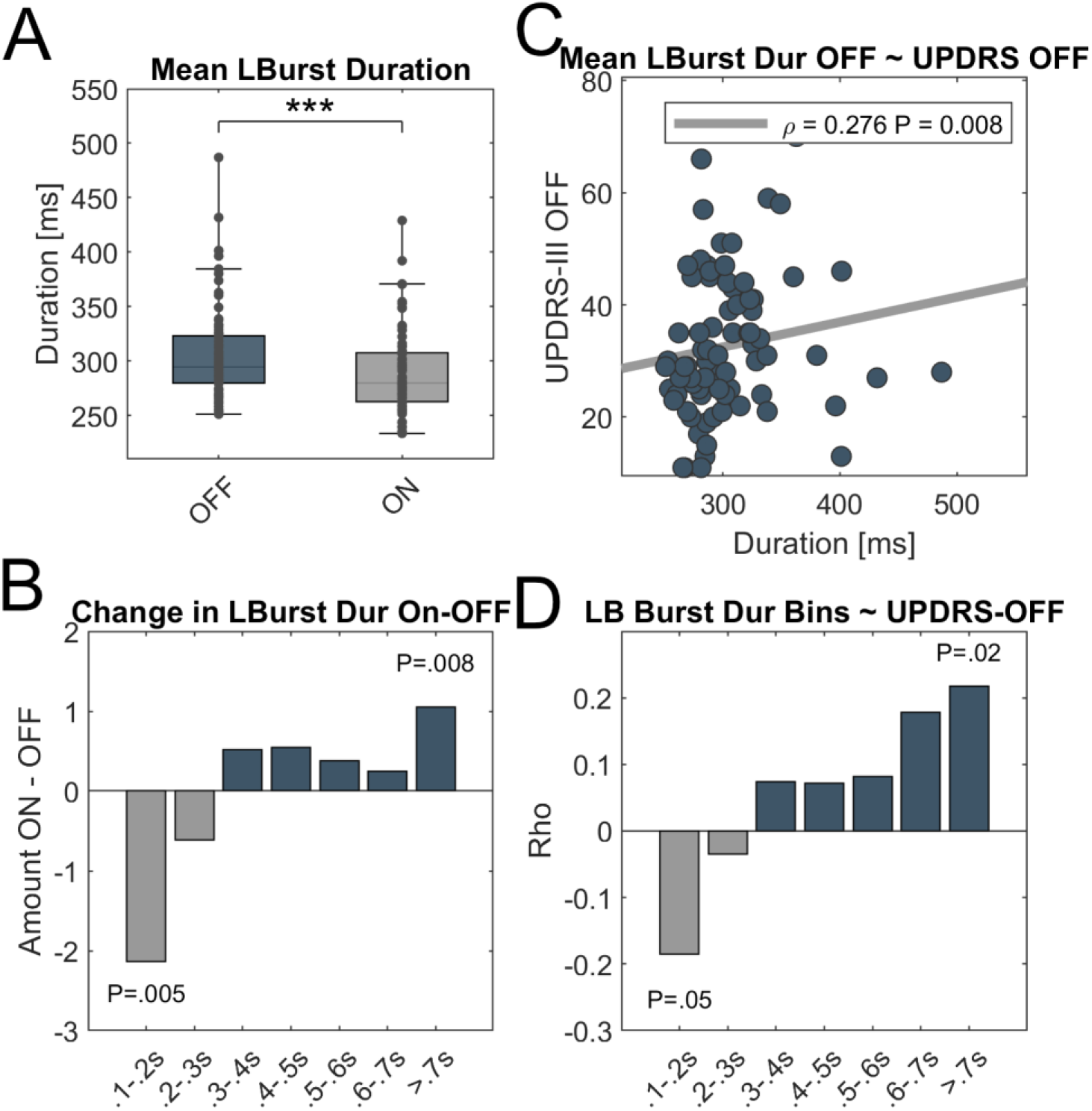
Low beta dynamics using the common threshold method based on the 75th percentile of the amplitude distribution. (A) There is a significant shortening of averaged low beta burst duration in the ON-medication state (P<0.001). (B) The increasing burst duration OFF-medication relies on a shift from very short bursts (100-200 ms) to bursts with prolonged duration exceeding 700 ms. Shown is the percentage change from the ON- to the OFF-medication state. (C) Motor impairment shows a significant positive correlation with the averaged LB burst duration. (D) When tested separately, the amount of very long bursts in the OFF-medication state correlates positively with symptom severity (Rho=0.22, P=0.02), while the amount of very short LB bursts show a trend towards a negative correlation with motor impairment. However, both results do not remain significant after FDR-correction. In boxplots, central marks indicate the median and edges the 25th and 75th percentiles of the distribution. *** p<0.001.

## Acknowledgements

We would like to thank all patients for their participation.

## Author Contributions

**Roxanne Lofredi:** conception and design of the study, acquisition and analysis of data and drafting a significant portion of the manuscript or figures.

**Liana Okudzhava:** analysis of data.

**Friederike Irmen:** acquisition and analysis of data.

**Christof Brücke:** acquisition of data.

**Julius Hübl:** acquisition of data.

**Joachim K. Krauss:** acquisition of data.

**Gerd-Helge Schneider:** acquisition of data.

**Katharina Faust:** acquisition of data.

**Gerd-Helge Schneider:** acquisition of data.

**Wolf-Julian Neumann:** acquisition and analysis of data.

**Andrea A. Kühn:** conception and design of the study, drafting a significant portion of the manuscript.

