## Supplementary figures and images for "Subthalamic beta bursts correlate with dopamine-dependent motor symptoms in 106 Parkinson’s patients"

### Supp. Figure 1

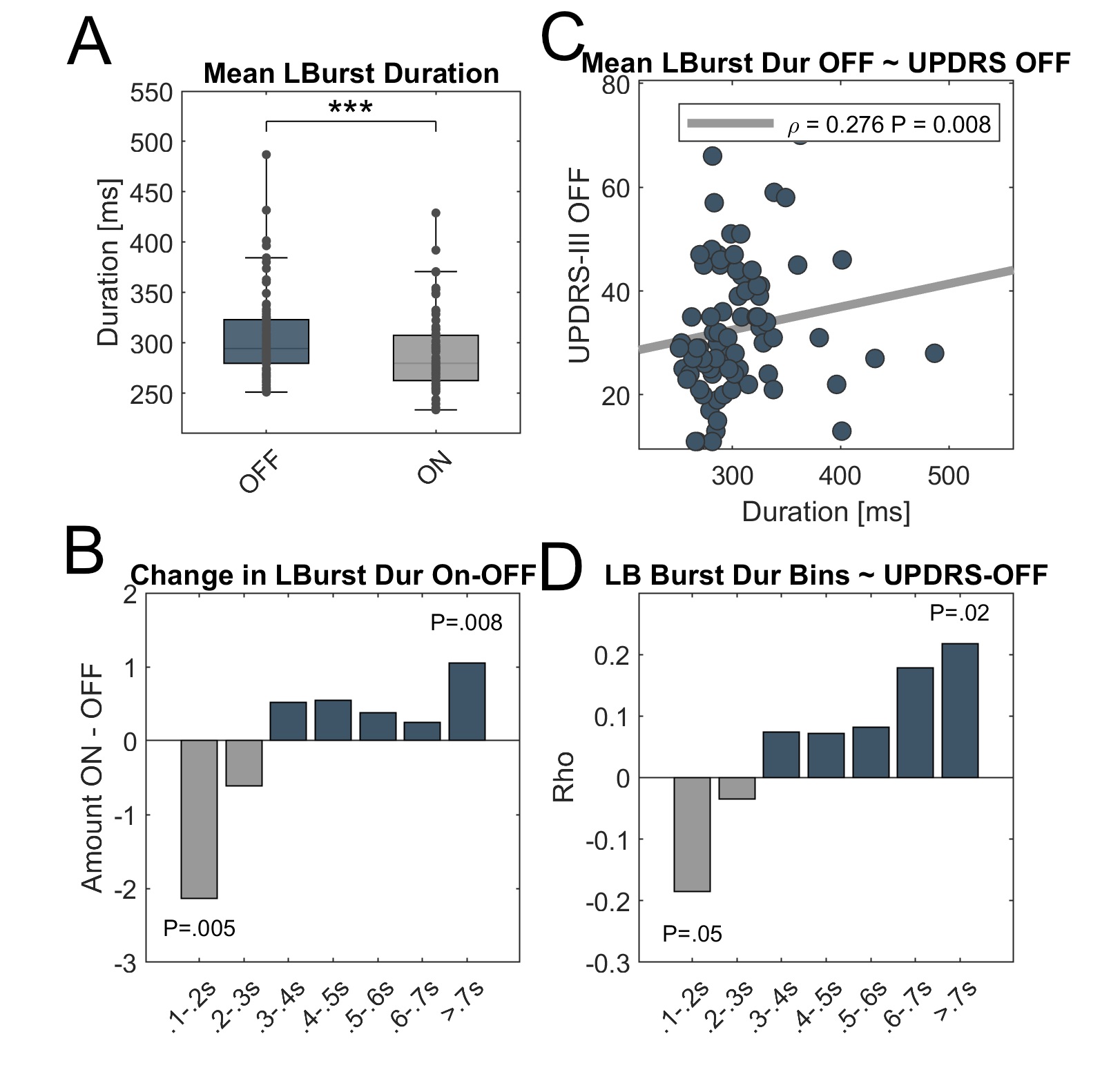
